# Cytocapsulae and cytocapsular tubes are golden targets for precise, effective and efficient cancer diagnosis and therapy in 307 kinds of liquid and solid cancers

**DOI:** 10.1101/2022.02.03.478975

**Authors:** Tingfang Yi, Gerhard Wagner

**Author notes:** Corresponding author: Dr. Tingfang Yi, Cytocapsula Research Institute, 245 First Street, Cambridge, MA, 02142 USA, or at.

## Abstract

Cancer is a leading cause of human lethality worldwide. A cancer cell is not only the cytoplasm and nucleus enclosed in the cell membrane. A malignant tumor is not only a mass of cancer cells. Cancer in a patient is not only isolated tumors distributed in tissues and organs. The comprehensive understanding of the complete structural compartments and their biological functions of cancer cells, malignant tumors and tumor systems in patients *in vivo* is an indispensable prerequisite for effective drug development and efficient clinical cancer pharmacotherapies. However, there remains an open question for a long time: what are the complete structural compartments of cancer cells, malignant tumors and tumor systems in the patient body? Here, we comprehensively and systematically investigated the complete structural compartments of cancer cells, malignant tumors and cytocapsular tube (CCT) network-tumor system (CNTS) of 307 types/subtypes of cancers in 33 kinds of tissues and organs by biochemistry and molecular biology assays with 9,811 clinical cancer tissue specimens from 9,637 cancer patients and >110,000 immunohistochemistry (IHC) fluorescence staining images. We discovered the complete structural compartments of cancer cells, malignant tumors and CNTSs in 307 kinds of cancers, elucidated mechanisms of cytocapsular growth and CCT elongation, and identified the comprehensive procedures of cancer evolution in patients. This study paves avenues for curing all kinds of cancers.

## Introduction

Mankind has spent extremely intensive efforts in the war against cancer in the last 5,000 years since first written records dated back to 3500–3100 BC^1^. By now, approximately 5 million scientific articles are documented in PubMed since 1783. US FDA approved about 650 cancer drugs since1949, and thousands of kinds of cancer therapies are applied in the world every day. The global cost of cancer was approximately $2.5 trillion in 2010 alone^2^. However, it is estimated that about 10 million cancer deaths and around 20 million new cancer cases occur in 2020 alone worldwide^3^. The marginal cancer pharmacotherapy outcomes suggest that the complete structural compartments and major mechanisms of the nature of solid and liquid cancers are still beyond our understandings^4-7^.

Why did most cancers remain uncurable despite the 5 million documented scientific articles, more than 2000 cancer drugs approved by all countries, and around $2.5 trillion annual expenses? The major facts below may provide comprehensive explanations to this open question: 1) the conventional understanding of structural compartments of cancer cells, malignant tumors and tumor systems in cancer patients is widely incomplete; lacking the understanding of the major structural compartments may prevent all conventional cancer therapies from effectively hitting the major targets and therefore only yield marginal outcomes; 2) the fragmental understandings of the complete structural compartments and mechanisms of cancer cells, malignant tumors and tumor systems unfortunately lead to directional mistakes, indecisive cancer prognosis and diagnosis, misdiagnosis, false negative diagnosis, ineffective and inefficient drugs, and all kinds of therapies with marginal outcomes; 3) the low value of conventional cancer cell assays is based on incomplete cancer cells that do not contain cytocapsular tubes *in vitro*. This also applies to all kinds of low clinical prediction value of animal assays (such as cancer cell derived xenograft (CDX) and patient cancer cell derived xenograft (PDX)) which lack the major structural compartments in clinical malignant tumors; 4) all current cancer researches only focus on around 10% of genes and proteins, and the leave out 90% of the human genome and proteome are still hidden in the “black box” due to many limitations and restrictions^8^; 5) the majority of the 20,000 coding proteins, tens of thousands of protein types, millions of protein species, and billions of protein molecules, the hundreds of thousands of non-coding genes (“junk genes”) are largely unknown; 6) the non-essential classification of cancers into hundreds of or even thousands of subtypes based on fragmental, marginal, incomplete and unstable molecular markers (genes/proteins), non-inherent characters of cell morphology in 2D cell culture (in which CCTs are absent), leading to imprecise cancer diagnosis and poor therapy outcomes^9-14^; 7) there is a large volume of cancer drug requests from 90 millions diagnosed cancer patients and from approximate 1.03 billion suspicious cancer patients every year worldwide, requiring long-term (10-12 years/cancer drug) and expensive investment (average $2.7 billion/cancer drug) in cancer drug discovery and development. This forces all countries to lower their approval standards to let some drugs go to the market, even though they only show unstable and limited benefits and are far from curing cancers. Among the 7 above facts, the first 3 items are the essential causes that lead to the current situations of poor outcomes of cancer therapies.

From 2014 to 2017, we discovered a new organelle, dubbed cytocapsular tube (CCT), which is a compartment lined by a second membrane outside of and distinct from the cell membrane^15^. While the mechanisms that lead to generation of the CCTs are still largely unclear, we found that CCTs conduct migration of malignant cancer cells. Cytocapsular tubes are generated by single cancer cell by shedding of, bi-layered lipid biomembrane outside of the plasma membrane while wrapping the originator cell inside, allowing other cancer cells to join inside, and thus providing membrane-enclosed freeways for cancer cell migration and dissemination^15^. From 2018 to 2021, we found that CCTs are exclusively present in 202 kinds of clinical solid cancer cells, but absent in normal tissue cells or benign tumor cells. CCTs interconnect and form superlarge CCT networks in patient tissues. CCTs and networks are universally present in the original cancer niche, in paracancer tissues (conventionally names “normal adjacent tissues, NAT”), and metastatic tumors. We established that CCTs dominate cancer metastasis physical pathways in 202 kinds of solid cancers^16^. Furthermore, from 2020 to 2022, we identified that: 1) cytocapsular tubes and networks function as physical superdefence freeway systems conducting conventional cancer drug pan-resistant tumor metastasis (cdp-rtm); 2) conventional animal tests of CDX and PDX cancer cell masses do not generate CCTs and networks and therefore have low clinical prediction value; thus, we created the CCT xenograft method, CCTX, which exhibits CCTs and networks in the xenografted tumors for highly efficient cancer drug screening and selection; 3) CCT network-tumor system (CNTS) is an integrated physical target for the highly effective and efficient pharmacotherapy of solid cancers; and 4) identified that organ colonized cancer cell masses (CDX or PDX, should not be considered as tumors) do not engender CCTs and networks and have low clinical prediction value^17-19^. The above-mentioned studies on CCTs systematically demonstrated that the previously unknown and invisible CCTs and their networks play key roles in broad-spectrum processes in cancer cell development, tumor formation, cancer metastasis. They are essential for growth and formation of metastatic secondary tumors, CNTS formation, conventional cancer drug pan-resistant tumor metastasis (cdp-rtm), cancer diagnosis, and requiring radio-, immune-, chemo-, and physical-therapies. However, whether cytocapsulae (CC) and CCTs are involved in the complete structural compartments of cancer cells, malignant tumors, and cytocapsular tube network-tumor systems of solid and liquid cancers is unknown.

It is well known that the cell membrane is the boundary of human cells. Recently, we discovered that CC and CCTs are temporospatial organelles and boundaries of cancer cells *in vivo*, which significantly expands our understandings of cancer cells in structure, compartments, biological functions, cancer evolution, diagnosis, and therapies. Evolutionally, cytocapsular tube membrane outside of the cell membrane of single cells is not unprecedented. There are three kinds of alive ancient cells that have bi-lipid layered biomembranes outside of the cell membrane: the gram-negative bacteria superfamily, chloroplasts, and mitochondria have appeared at diverse times billion years ago^20-22^. A gram-negative bacterium of photosynthetic bacteria evolved between 2.3-2.7 billion years ago^22^. Chloroplast arose from endosymbiosis of free-living cyanobacteria by eukaryotic cells ∼2 billion years ago. Mitochondria arose from the endosymbiosis, retention, and integration of a free-living ancient bacterium into a host cell more than 1.45 billion years ago^20^. It is projected that in the ancient ocean periods from 1.45 billion to 2.7 billion years, the double bio-membrane structure provided multiple advantages: extra-protection shielding stressful or extreme extracellular environments outside, creating more stable conditions within the cytoplasm membrane for bio-molecule interactions, evolution, and survival of cells under stressful microenvironments. It is well-known that uncontrolled cancer cell proliferation leads to stressful or extreme extracellular environments in place, including: nutrient deprivation, hypoxia, low pH with enhanced concentrations of lactic acids and other kinds of acids, increased toxicity with elevated metabolic wastes, lack of sufficient growth factors, increased competition between neighbor cancer cells, viscoelastic extracellular matrices, and so on. Based on our reliable induction of cytocapsular tube generation in 25 kinds of human cancer cells *in vitro*, it is known that the stressful ECM with appropriate biochemical, biophysical and biomechanical signals stimulate cancer cells to reactivate the evolutionally ancient capacities of generation of cytocapsular membranes outside of cancer cell membranes in patients *in vivo*. The shared parts of the structure of the biomembrane outside the cell membrane, and partially-shared biological functions with gram-negative bacteria, chloroplast, mitochondria, and cytocapsular tube membrane of cancer cells, provide hints that the generation of cytocapsular membranes of cancer cells may be a temporospatial reverse evolution process of the outer membrane. These observations may open a novel and unique view to understand the development of cancers at the evolution and devolution levels.

## Results

### Structural compartments of cancer cells and cancer cell masses, and mechanisms of cytocapsular growth

In order to investigate whether compact cancer cell masses generate superlarge cytocapsulae (CC) and mechanisms of how CC grow up, we first explored cancer cell masses on the 3D Matrigel matrix (MM) *in vitro* mimicking the microenvironments *in vivo* with immunohistochemistry (IHC) staining with anti-CM-01 (CC marker protein) and anti-gamma actin antibodies as well as fluorescence microscope analyses. Initially, single pancreas cancer Bxpc3 cells generate small CC outside of the single Bxpc3 cell. Bxpc3 cells continue to proliferate in the small CCs and form compact cancer cell masses (CMs) (**Fig. 1A**). Bxpc3 cells continue to divide and CMs grow up. CM cytocapsula sizes increase along the CM growth (**Fig. 1A**). Subsequently, all the Bxpc3 cells grow up into spherical CMs (these spherical CMs should not be considered tumorspheres as they lack of CCTs and networks inside), and all the CMs consistently generate superlarge CCs outside, wrapping the spherical CMs (**Fig. 1A**). The superlarge CCs enclose the CMs in spherical or irregular-shaped morphologies (**Fig. 1A**). *In vitro*, Bxpc3 cells of the CM inside the superlarge cytocapsulae do not engender their own CCs (**Fig. 1A**). Interestingly, Bxpc3 cells can disassemble the CMs, ecellulate, and evict outside of the big cytocapsulae of the CMs and leave ecellulated, concaved, membrane-enclosed discs without holes in the discs (**Fig. 1A**). Sometimes, multiple Bxpc3 cells ecellulate at the same time and leave an open hole (not reclosed) in the ecellulated CM cytocapsulae in a thick and short fragment/tail (**Fig. 1A**). These observations demonstrated that: 1) compact cancer cell masses can generate superlarge CC outside of CMs, 2) these CM cytocapsulae enclose CM inside, and shield cancer drugs outside and reduce the drug access to cancer cells inside the CC, 3) cancer cells of CM can disassemble and leave CM individually or collectively via CCTs or ecellulation (**Fig. 1A**).

**Fig. 1.**
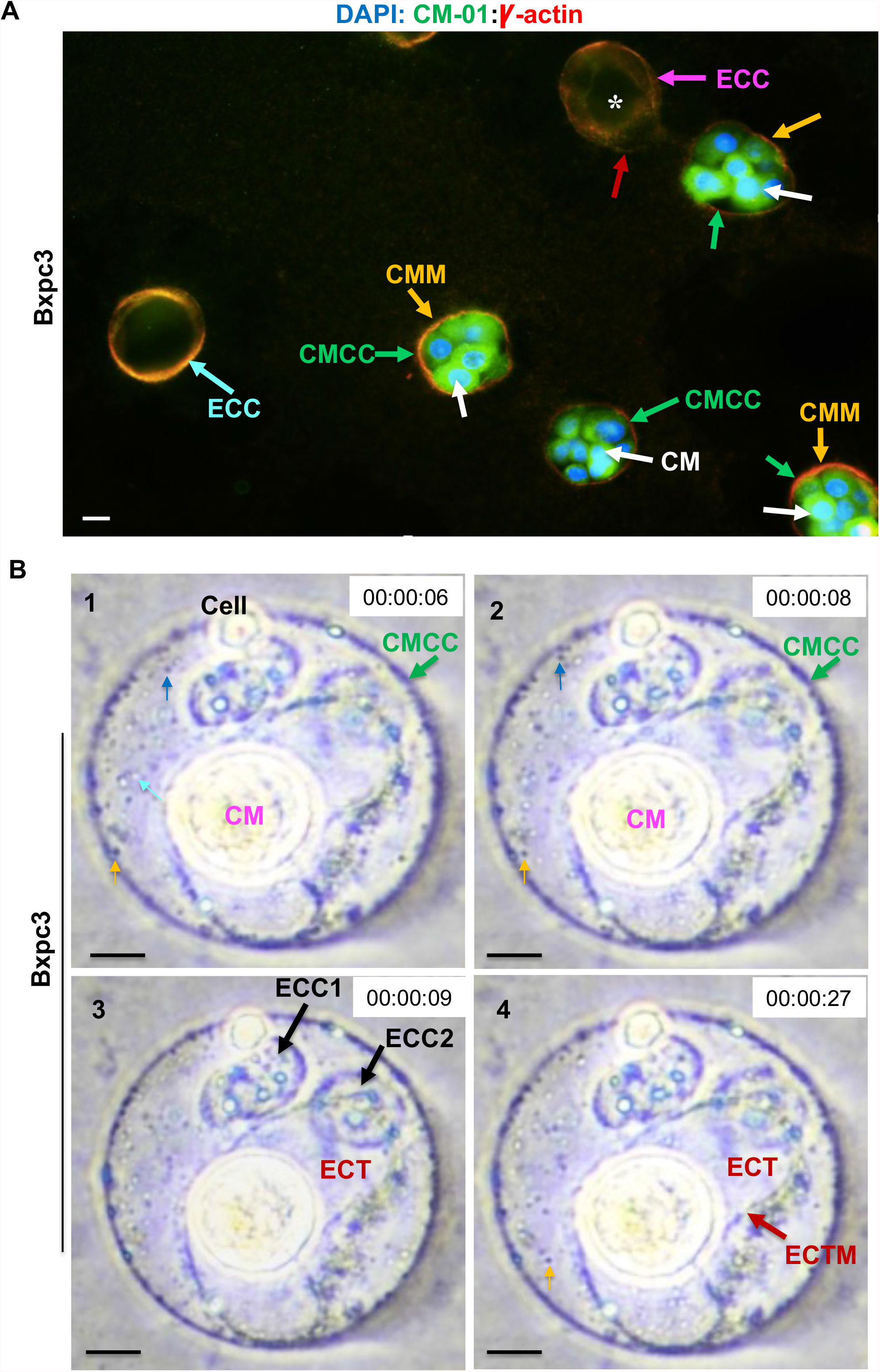
Characterization of complete compartments of clinical cancer cells. **(A)** A representative fluorescence microscope image shows that pancreas cancer Bxpc3 cell masses generate superlarge cytocapsulae outside and wrap cancer cell mass (CM) inside on the thick Matrigel matrix in 6-well plate *in vitro*. The immunohistochemistry-fluorescence staining was performed with anti-CM-01 (protein code) and anti-gamma actin antibodies and DAPI. Usually, all cancer cells in CMs individually ecellulate and leave big, round, reclosed, reunified, and ring-shaped double concaved discs without holes (in the left in the image, gray blue arrow). Sometimes, the CMs can massively ecellulate and leave a big open hole (labeled with white asteroid star) without reclosed or reunified (at the top in the image, purple arrow). The individual cells in the CMs within the superlarge cytocapsulae (CMCC, green arrows) do not generate their small CCs in this study. The cancer cell mass (CM, white arrows), superlarge cancer cell mass cytocapsulae (CMCC, green arrows), cell mass cytocapsulae membrane (CMM, orange arrows), ecellulated cytocapsulae (ECC, purple or gray blue arrows) without CMs/cells inside, and the edge of cytocapsulae membrane of the open hole of ECCs (red arrow) are shown. Scale bar, 10μm. **(B)** Representative multiple high resolution, bright field, time-lapse microscope images of Bxpc3 cell superlarge cancer cell mass cytocapsula growth with the fusion of thousands of cancer cell secreted, tiny, membrane enclosed vesicles. There is a Bxpc3 cancer cell mass (CM) in the middle, which has engendered a superlarge cytocapsula (CMCC, green arrow). In the CMCC lumen and beyond the cell mass, there are thousands of tiny, membrane-enclosed vesicles (appear as focused tiny black/dark gray dots (blue or orange arrows) or unfocused golden tiny dots (cyan arrow) in the CMCC lumen in the image due to different layers in the big CMCC bubble lumen and the always moving of the tiny vesicles; thin orange or blue arrows two of them) randomly moving outside of the CM. In the panels 1 and 2, the blue arrows show a tiny vesicle move to close to the CMCC membrane in 2 seconds. In the panels of 1, 2 and 4, the orange arrows show a tiny vesicle randomly move forward to the center of the CMCC lumen. In the panel 3, there are two ecellulated cytocapsulae (ECC, ECC1 and ECC2, thick black arrows) in the cancer cell mass cytocapsula lumen; the cell individuals evicted and left the CMCC and floated away. In the panel 4, the thick red arrow shows an ecellulated cytocapsular tube (ECT) in a short, big, and L-shaped morphology, with the cell ecellulated and left the cytocapsular tube (CCT) and CMCC. The thick red arrow shows the ECT membrane (ECTM). The ecellulated cytocapsulae and cytocapsular tube in the CMCC show that cancer cells can go through multiple layers of cytocapsular membranes during CC/CT ecellulation. Time in the movie is shown. Scale bar, 10μm.

Next, we explored how CMs’ cytocapsulae grow up. Using high resolution bright light time-lapse microscope, we take high resolution of alive movies of pancreas cancer Bxpc3 CM inside CC. Interestingly, there are hundreds or thousands of tiny (<0.2μm in diameter/width), membrane-enclosed, and different kinds of vesicles inside the CM cytocapsular lumens, but outside of the cancer cells masses and the CCs of single cancer cells (**Fig. 1B**). These tiny membrane-enclosed vesicles are in constant and random movement in irregular ways within the CM cytocapsulae lumens. These tiny membrane-enclosed vesicles are secreted from the cells of the CM (**Figs.1B and S1A**) and most of them later reach and attach the inner side of the membranes the CM superlarge CC. Then, these large quantities tiny membrane-enclosed vesicles fuse into the CM cytocapsulae membranes, through which the CM cytocapsulae enlarge and grow up (**Fig. 1 and S1A**). These observations demonstrated that single cancer cells secrete large quantities of tiny (<0.2μm in diameter/width) membrane-enclosed vesicles to form cytocapsulae outside of the single cancer cells, and to fuse into the small cytocapsular membrane, increase CC membranes, and subsequently generate larger cytocapsulae. There are multiple different kinds of tiny membrane-enclosed vesicles secreted by the cancer cells in the CC or CCT lumens, in which molecular components and biological functions need more study to elucidate.

Sometimes, single foreign cancer cells with their own small CC can enter the CCs of CMs without CC degradation. After entering the CM cytocapsulae lumens, the single foreign cancer cells perform double-ecellulation from their own small CC (or short CCTs if migrate and generate short CCTs inside) and the CC of other cancer cell masses, leaving two ecellulated small CC and a short CCT in the lumens of CM cytocapsulae (**Fig. 1B**). Sometimes, the superlarge CCs of CMs generate multiple spike-like structures in the spherical lumen linking the cancer cell masses and the CC membranes, which facilitate to support and maintain the superlarge CC (**Fig. S1A**).

Next, we investigated whether cancer cell masses in cancer patient malignant tumors *in vivo* generate superlarge cytocapsulae. We examined breast cancer cell masses in the clinical breast carcinoma specimens. We found that the compact breast cancer cell masses in breast carcinoma (malignant tumors) generate superlarge cytocapsulae localized outside of the huge breast cancer cell masses, wrapping the huge cancer cell masses (**Fig. 2A**). Most single breast cancer cells inside the superlarge CC of the huge cancer cell masses do not engender small CC of their own, which is consistent with the phenomena *in vitro* (**Figs. 1A and 2A**).

**Fig. 2.**
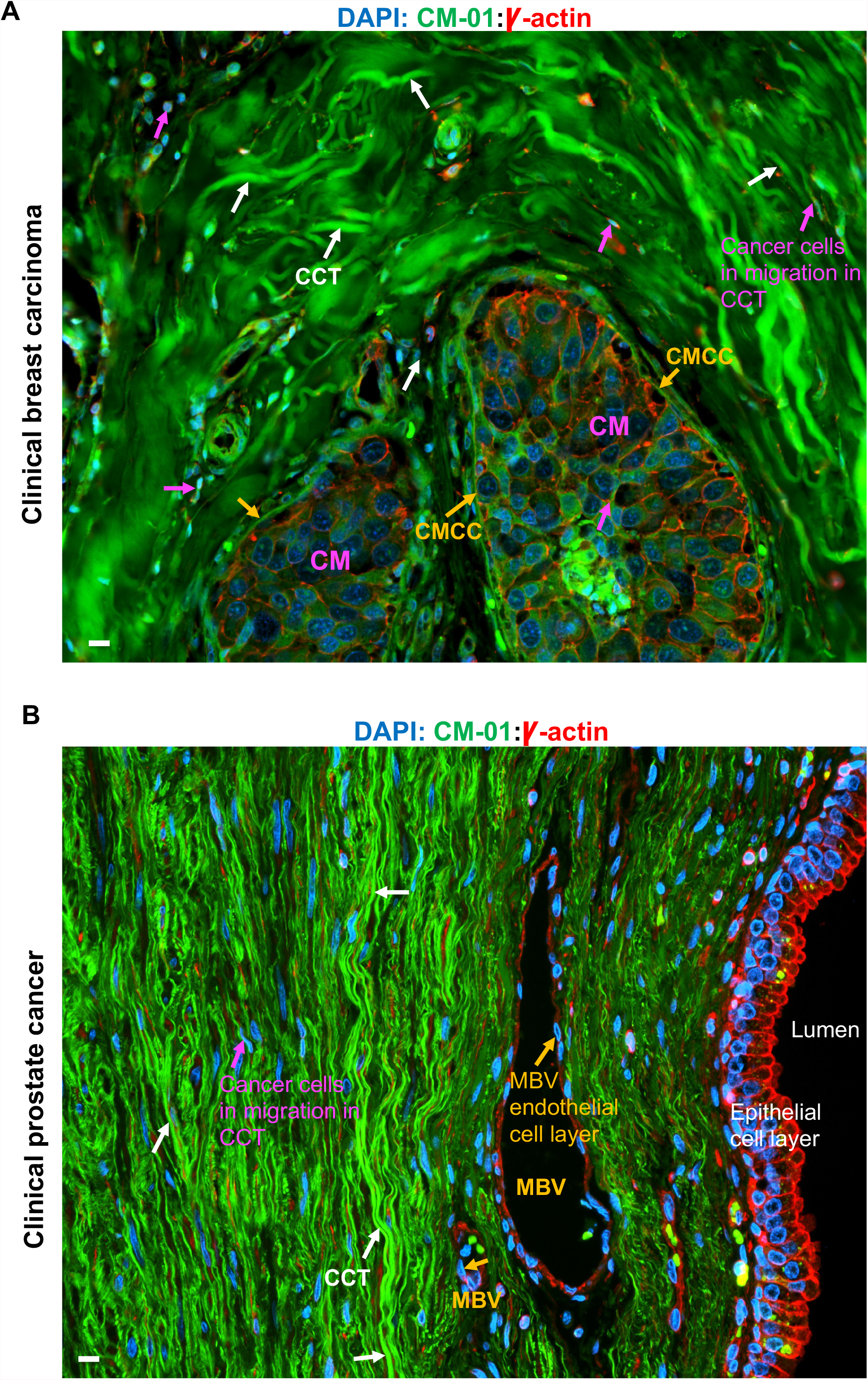
Characterization of complete compartments in clinical primary malignant tumors. **(A)** A representative fluorescence microscope image of clinical breast carcinoma area with multiple compact and dense cancer cell masses (CMs), cytocapsular tubes (CCTs, white arrows), and cancer cells in migration in CCTs. The superlarge and very thin membranes of cancer cell masses cytocapsulae (CMCC, orange arrows), and cancer cells in migration in CCTs (purple arrows) are shown. Scale bar, 10μm. **(B)** A representative fluorescence microscope image of clinical prostate malignant tumor area with large quantities of cytocapsular tubes (CCTs, white arrows), micro blood vessel (MBV), integrated and tightly assembled MBV endothelial cell layer (orange arrow) without be invaded by CCTs at cancer Stage IIc, prostate cancer cell in migration in CCTs (purple arrows). Scale bar, 10μm.

**Fig. 3.**
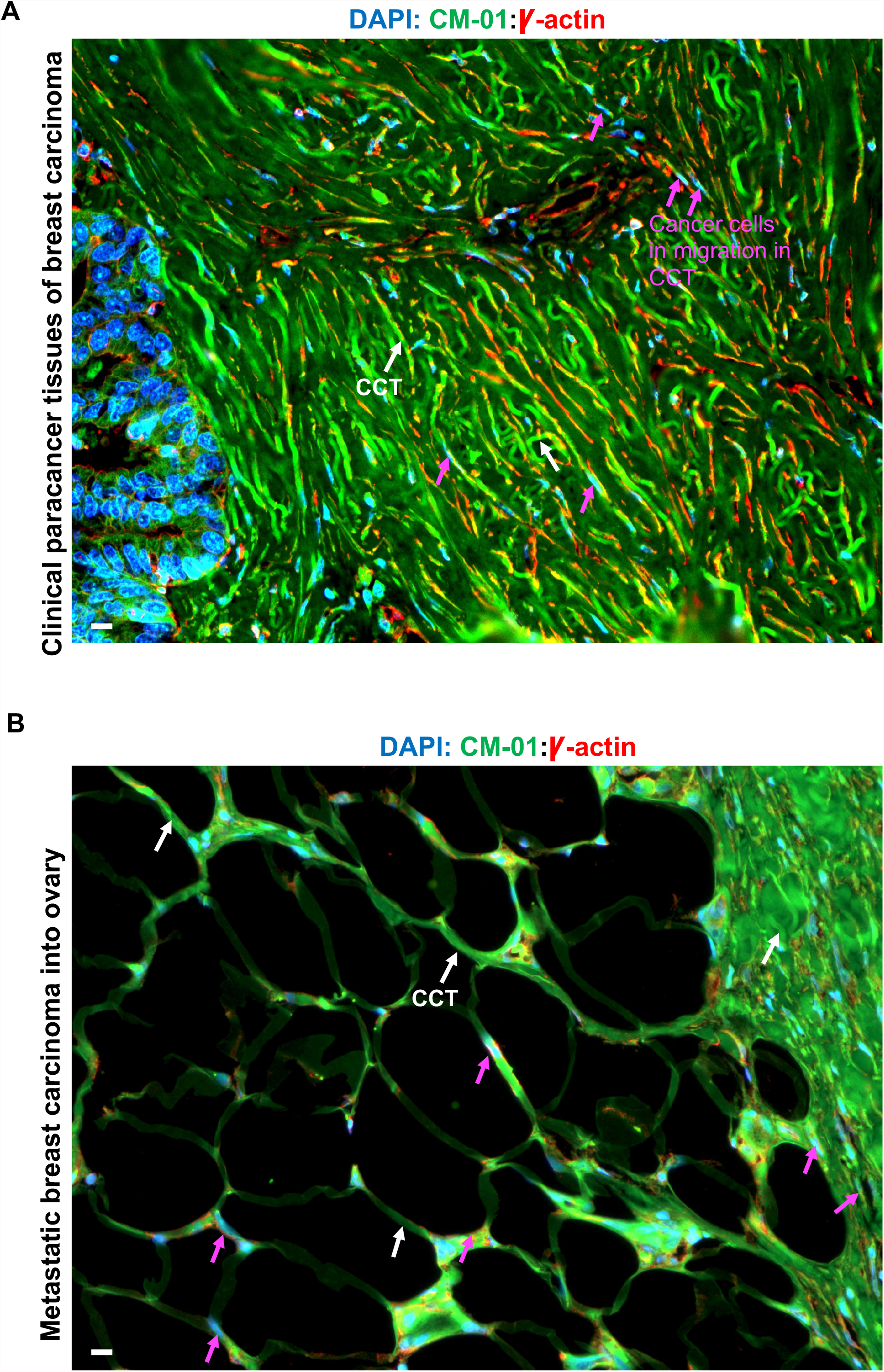
Characterization of complete compartments in cytocapsular tube (CCT) network-tumor systems of primary and metastatic secondary tumors in cancer patients. **(A)** A representative fluorescence microscope image of paracancer tissue area of breast cancer. There are millions of breast cancer cytocapsular tubes (CCTs, white arrows) piled up and form compact CCT masses in the paracancer tissues (previously called “normal adjacent tissues, NAT”, which are false negative normal tissues). Large quantities of breast cancer cells are in migration in breast cancer CCTs at the paracancer tissues and move forward to far distance destinations. Scale bar, 10μm. **(B)** A representative fluorescence microscope image of metastatic breast cancer in ovary in cancer patient. There are superlarge, 3D, sparse, breast cancer CCT networks and highly compact CCT networks (right side) in the same area in metastatic secondary breast tumor in ovary. Many breast cancer CCTs (white arrows) interconnect and form superlarge CCT networks and many breast cancer cells are in migration in loose or dense CCT networks (purple arrows) in breast tumors in the ovary. Scale bar, 10μm.

In short, the above data demonstrated that the complete structural compartments of **single cancer cells** include 4 major compartments in the order from outside to inner side: 1) cytocapsulae or cytocapsular tubes outside of the cell membrane, 2) cytocapsular lumen fluids containing many tiny, membrane-enclosed vesicles, 3) cell membrane and the cytoplasm, and 4) nuclear membrane and the nucleus (**Fig. S1B**). The compartment items of 1) and 2) have previously not been seen and were unknown. Cytocapsulae and cytocapsular tubes shield stressful environmental stimuli outside, protect cancer cells inside, provide physical, membrane-enclosed, superdenfence freeways for cancer cell dissemination, significantly increase the survival of cancer cells in stressful microenvironments. The tiny membrane-enclosed vesicles are secreted by the single cancer cells and are building blocks of the cytocapsulae and cytocapsular tubes. The tiny membrane-enclosed vesicles fuse to cytocapsulae and cytocapsular tubes, increase CC/CCT membrane surface areas, and drive the growth and elongation of cytocapsulae and cytocapsular tubes.

The complete structural compartments of **cancer cell masses** in cancer patients include 3 major structural compartments in the order from outside to inner side: 1) superlarge cytocapsular membranes wrapping cancer cell masses, 2) the cytocapsular lumen fluids containing many tiny, membrane-enclosed vesicles, 3) the compact cancer cell mass in the CCs (**Fig. S1C**). The above items (1) and 2) have previously not been seen and were unknown. Cancer cells proliferate in the CC; cancer cells secret a lot of tiny, membrane-enclosed vesicles, which fuse into the CC membranes, and increase the CC membranes, and enlarge the CC sizes. CC sizes increase along the cancer cell proliferation and cancer cell masses grow up. The superlarge CC membranes enclose CMs inside, constantly shield stressful environmental stimuli outside, protect cancer cell masses inside. The cancer cell mass CC linked CCTs provide physical, membrane-enclosed, and superdenfence freeways for cancer cell dissemination, significantly increase the survival of cancer cells in the CM in stressful microenvironments.

### Complete structural compartments of malignant tumors of 307 kinds of cancers in patients

Next, we investigated the complete structural compartments of malignant tumors in patients *in vivo*. Using fluorescence microscope with anti-CM-01 and anti-gamma-actin antibodies, we comprehensively analyzed >110,000 immunohistochemistry (IHC) fluorescence images of 9,811 clinical malignant tumors (both original niche and metastatic secondary tumors) from 9,637 cancer patients of 5 countries, covering 3 continents: America, Europe and Asia. These randomly tested 9,811 clinical malignant tumor (cancer) tissues exhibit extreme heterogenicity and diversity in tissue texture, composition, morphologies, compact cancer cell masses, cytocapsular tubes in all kinds stage in CCT life cycles, CCT morphologies, superstructures, CCT networks, CCT density, cell types, micro blood vessels, veins, cell density, cell status (proliferation, apoptosis, necroptosis), cell types, cell morphologies, cell sizes, and molecular protein abundances. By integration all of the comprehensive and systematic information, we set up the complete structural compartments of malignant tumors *in vivo*.

The complete structural compartments of **malignant tumors** in patients include 5 major structural compartments: 1) superlarge cytocapsulae enclosed compact cancer cell masses (CMs): in these CMs most single cancer cells initially do not generate their own CC; the outer layer single cancer cells can invade into the tightly contacted CCTs by penetrating the superlarge CC membrane and the CCT membrane of other cancer cells; later, single cancer cells in the CM within superlarge cytocapsulae can generate their own CC and CCTs, and these CCTs can invade outside of the superlarge CCs and leave CMs, relocate to far distance destinations (**Fig. 2A**); 2) CCTs and CCT networks in all kinds of stages in CCT life cycle: CCTs broadly interconnect and form 3D, complex and large networks in all kinds of morphologies, textures, structures, and densities; these CCTs and networks potently invade, extend to and pass through all kinds of paracancer tissues, lymph nodes (in local, neighboring tissue, and far distance), dense or loose tissues, normal or extremely high cell densities (**Fig. 2, Table 1**); 3) CCT network-cancer cell masses systems in the original or metastatic secondary malignant tumors: cancer cells migrate in the CCTs and networks, and assembly or disassembly between the CMs in the malignant tumors (or cancer blocks) (**Fig. 2, Table 1**); 4) CCTs wind outside of blood and lymph vessels: before cancer Stage IIC, no CCTs invade into micro blood vessels and no solid cancer cells/fragments are released into the circulation systems, but millions or hundreds of millions of cancer cells already migrate and disseminate to neighbor or far distance sites through the hundreds of thousands or millions of CCTs; after Stage III, <1 in 1million CCTs invade into vessels and release cancer cells into the circulation (the resources of circulating tumor cell (CTC), and the base of ctDNA diagnosis, leading to misdiagnosis, false negative diagnosis for pre-stage, Stages I and IIc, and later stages); CCT networks are independent of the circulation systems, providing membrane-enclosed, superdefence, freeway systems for CCT-guided and protected metastasis; CCTs invade into micro blood vessels and lead to blood vessels leaky and even decomposition, leading to low blood pressure in the late stages; many kinds of blood based tests (such as colon blood plasma test in the feces) cannot diagnose of early solid cancer stages (Stages I and II), and will make indecisive diagnosis for the late stages III and IV (**Fig. 2, Table 1**); 5) mixed tissues and cells: they can be normal tissues, benign tumor cells, neoplasia cells, single cancer cells in their CC (in compact or loose tissues) (**Fig. 2, Table 1**). These malignant tumors can be in extremely diverse statuses: compact blocks (with increased cell density, and can be detected by CT-scan if >1mm in sizes), loose tissues sporadically distributed in the normal tissues (without increased cell density, will be invisible by all conventional diagnosis methods), highly dense CCTs and networks (without increased cell density, leading to false negative diagnosis, misdiagnosis, false positive effective cancer therapies), highly degraded CCTs and networks with many CCT cavities (without increased cell density, leading to leading to false negative diagnosis, misdiagnosis, false positive effective cancer therapies); many kinds of polyps in colon, stomach contain CCTs and networks, but conventional diagnosis methods cannot analyze them.

**Table 1.**
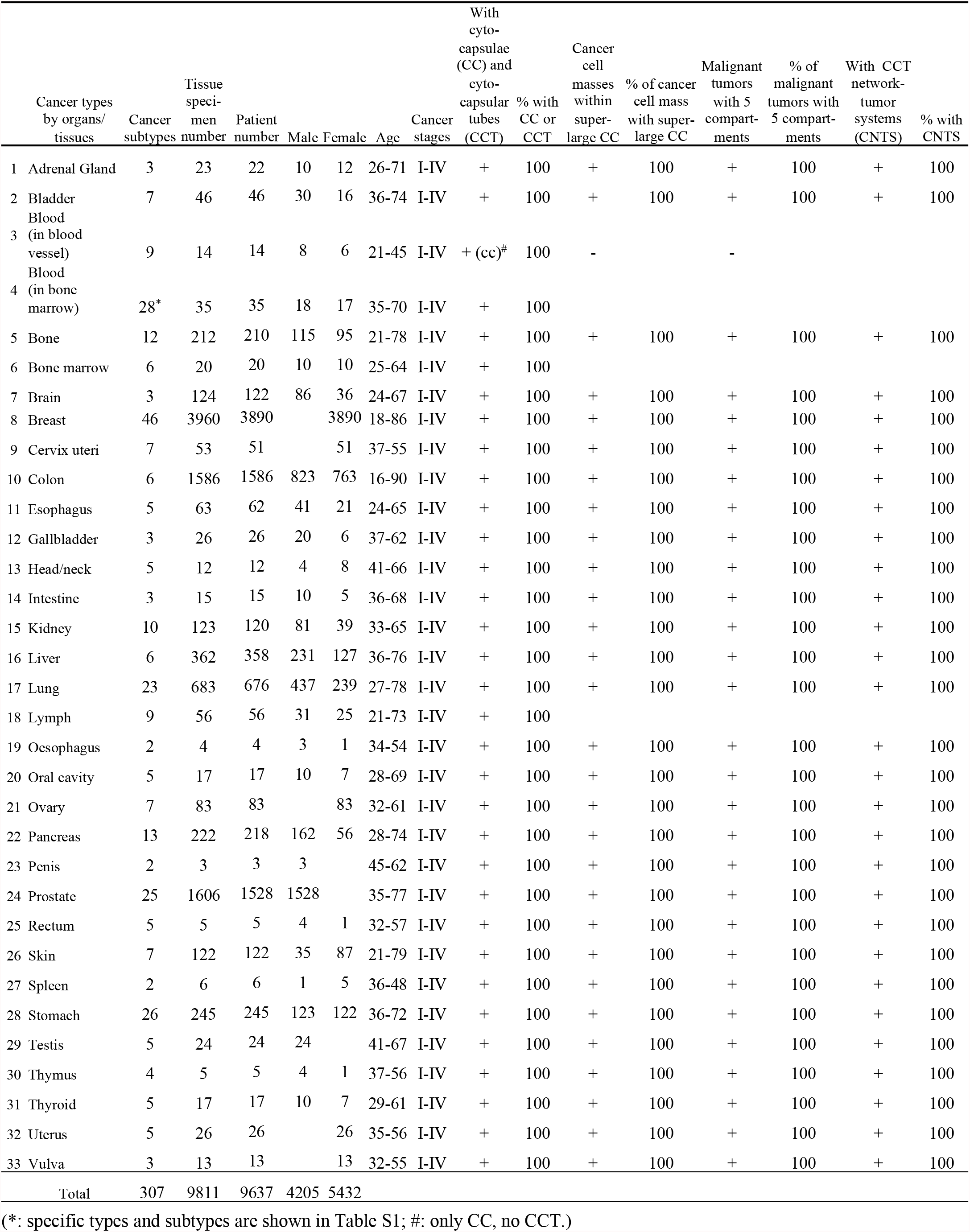
Characterization of complete compartments of cancer cells, malignant tumors, and CNTSs in 307 types (subtypes) cancers in 32 kinds of organs/tissue in human.

In short, the above described 5 complex compartments of clinical malignant tumors make it very difficult for conventional diagnosis methods to reach precise, quantitative and decisive diagnosis, and for conventional radio-, immune-, chemo-, and physical-therapies to achieve highly precise, effective and efficient therapy and satisfied outcomes. On the other hand, the malignant tumor associated cytocapsular membrane systems with multiple important biological functions in all stages of malignant tumor evolution, which makes cytocapsular membrane system a golden target for the precise cancer diagnosis, and for the effective and efficient cancer therapy.

These comprehensive observations demonstrated that the conventional tumor experimentations in animals (such as CDX and PDX models; these xenografts cannot be considered as tumors or clinical mimicking cancer cell masses) lack most of the 5 structural compartments of malignant tumor. The cancer cell masses in conventional tumor analyses in animals lack the superlarge cytocapsulae outside of the cell masses (**Fig. 2, Tables 1, S1-3**). These unbiased results explain why the conventional tumor analyses in animals have very low clinical prediction value (3%- 4%)^23-24^ because these experimental cancer cell masses lack the 5 structural compartments in malignant tumors in cancer patients, and the cancer drug candidates selected by these experiments will not have the capacities to inhibit malignant tumors in cancer therapy.

In summary, the newly discovered 5 structural compartments of malignant tumors in cancer patients significantly expand our understandings of malignant tumors, and open avenues for cancer diagnosis and cancer therapies.

### Structural compartments of cytocapsular tube network-tumor system (CNTS) in 307 kinds of cancers in patients

Next, we explored the structural compartments of cytocapsular tube network-tumor systems in 307 kinds of solid cancers. We comprehensively analyzed >110,000 IHC fluorescence images of 9,811 clinical malignant tumors (before, during and after radio-, immune-, chemo- and surgery-, and physical-therapy; primary niches, paracancer tissue, secondary tumors) from 9,637 cancer patients (95% of them are dead patients after several times of pharmacotherapies). These 9,811 clinical malignant tumors display extreme heterogenicity and diversity in cancer cell masses, tumors, CCTs, CCT networks, cell density, cancer cell in migration in CCTs.

By integrating all information of the analyses of the indicated >110,000 high quality and unbiased IHC fluorescence images, we comprehensively and systematically achieve a description of the 4 structural compartments of **CNTS**: 1) all tumors in the patient (small or big in sizes, detectable or undetectable by conventional diagnosis methods); 2) superlarge and 3D CCT networks from the original niches to all secondary tumors and along all the metastasis ways: CCT networks interconnect all primary and secondary tumors; the superlarge and 3D CCT networks dynamically and broadly interconnect all tumors (visible or invisible by conventional diagnosis methods); cancer cell freely assembly to form new tumors in CCT terminal, or disassemble from and leave a tumor and reach to other tumors, or generate CCTs in different direction and grow up into a new tumor at a different site; some CCTs degrade and cancer cells will generate new CCTs and networks; 3) CCTs invade into bone and bone marrow: cancer cells reach bone marrows via CCTs and make niches inside, which significantly increases cancer cell survival and capacities for cancer relapse; 4) CCTs invade through blood-brain barrier and reach the brain (**Fig.3, Tables 1, S1-3**).

Interestingly, in the investigated 11 kinds of blood cancers in the blood, these blood cancer cell individuals generate CC, but not CCTs, outside of the cell membrane (**Tables 1 and S1**). In the investigated 28 kinds of blood cancers in the bone marrow, all these blood cancer cells generate a lot of CCTs and CCT networks in bone marrow. The examined 28 kinds of blood cancer cells migrate in blood cancer cell CCTs in the viscos bone marrow (**Fig. 4 and Tables 1 and S1-3**). The investigated 9 kinds of lymphoma generate CCTs in the lymph nodes and in bone marrow (**Table 1**). These observations demonstrated that: 1) the blood cancer cells generate CC in the blood and engender CCTs and CCT networks in the bone marrow stage; 2) blood cancer cells in the bone marrow stage need to migrate in CCTs and then are released into blood vessels; 3) lymphoma cells engender CCTs and CCT networks in both lymph nodes and bone marrow, and lymphoma cells migrate in lymphoma cell CCTs and then are released into the circulation vessels.

**Fig. 4.**
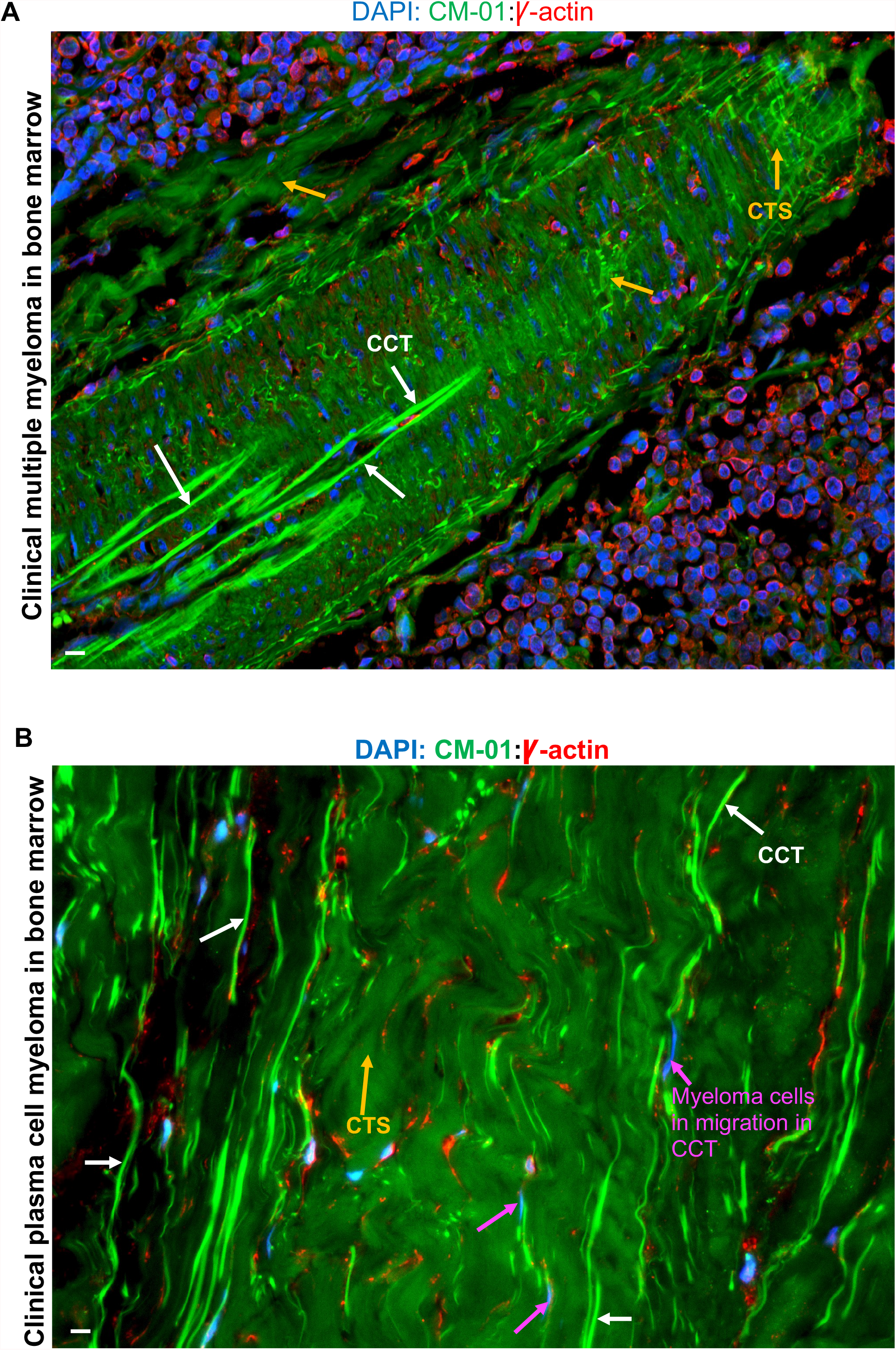
Characterization of complete compartments in clinical liquid cancers. **(A)** A representative fluorescence microscope image of clinical multiple myeloma in bone marrow. There are many long and winded multiple myeloma CCTs forming superlarge irregular shaped structures in the bone marrow. There are large quantities multiple myeloma CCTs degrade into thick, thin or very thin strands (CTS, orange arrows). Scale bar, 10μm. **(B)** A representative fluorescence microscope image of clinical plasma cell myeloma in bone marrow. There are many long plasma cell myeloma CCTs (white arrows) without degradation in the bone marrow. There are large quantities plasma cell myeloma CCTs degrade into thick, thin or very thin strands (CTS, orange arrows), with many in cloud-like status. Many plasma cell myeloma cells (purple arrows) are in migration in plasma cell myeloma CCTs and CCT networks in bone marrow. Scale bar, 10μm.

**Fig. 5.**
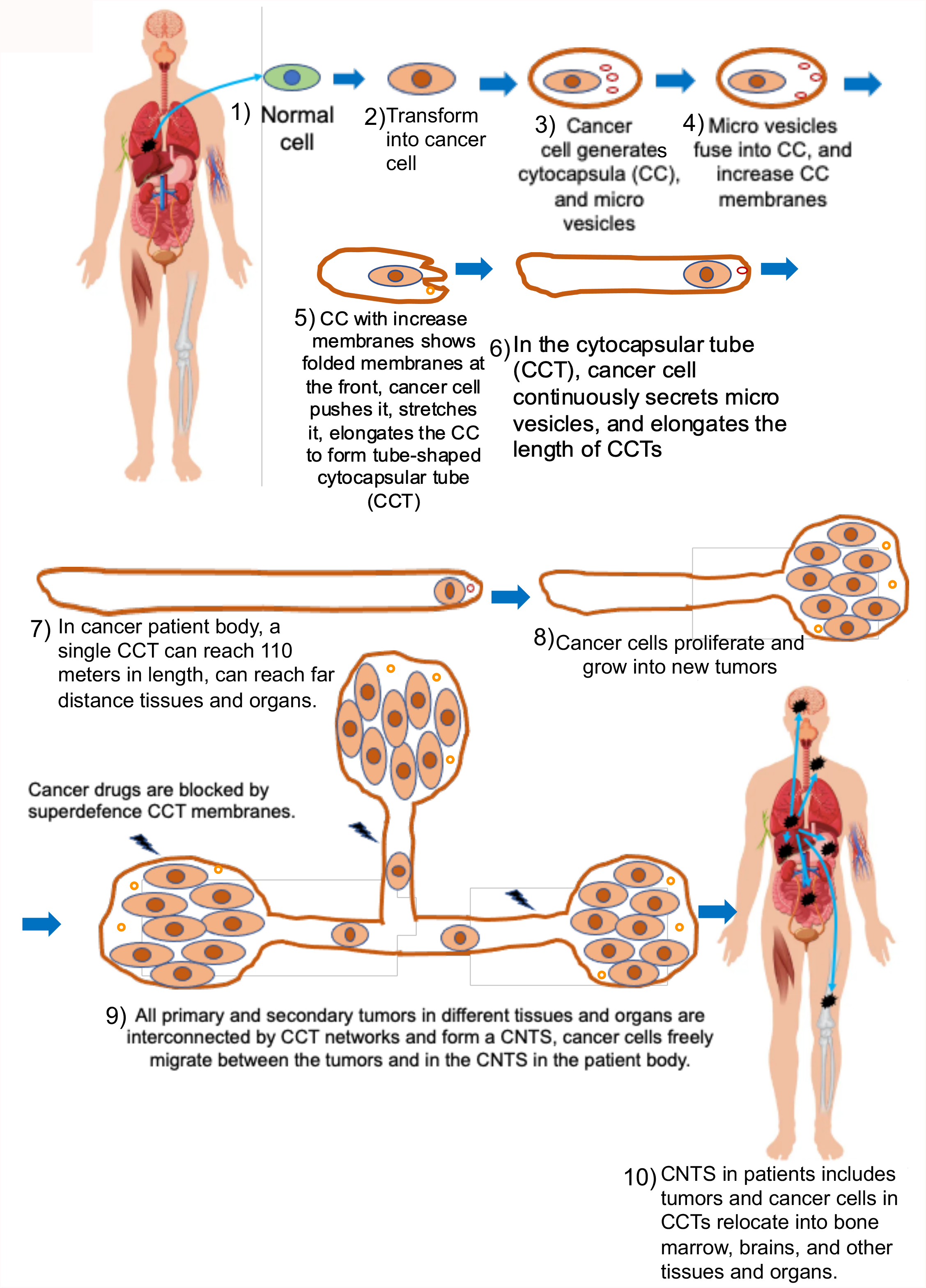
Diagram drawing of the complete procedure outlines of cancer evolution. The complete cancer evolution outline contains 10 major successive steps: 1. With radio-, chemical-, physical-, environmental stimuli and the accumulated gene mutations, a normal cell is transformed into a cancer cell. 2. Cancer cells have two major characters: uncontrolled proliferation, and cytocapsular tube conducted metastasis. Uncontrolled cell proliferation leads to local stressful environments with many stressors: nutrient deprivation, hypoxia, low pH (caused by elevated lactic acid and other acids), increased metabolic waste toxicity, insufficient growth factors, and increased competition between cancer cells, and so on. 3. Single cancer cell individuals are forced to generate cytocapsulae to wrap itself to physically separate and protect it from the stressful extracellular micro-environments. Single cancer cells constantly generate and release many tiny membrane-enclosed vesicles, 4. These tiny membrane-enclosed vesicles fuse into the small CC membrane, and increase the CC membrane size in area. 5. Single cancer cell push CC membrane, and stretch the increased and folded CCT membrane, elongate the CC in length, and create tube-shaped vessels, named as CCTs. At the same time, the single cancer cells secrete proteases via one (or some) kind(s) of the tiny membrane-enclosed vesicles acting as delivery shuttles. 6. The single cancer cells continuously secrete many tiny membrane-enclosed vesicles, fuse into the front CCT membrane, increase CCT membranes; secreted proteases outside of the CCT terminals and degrade ECM and making rooms for CCT elongation; single cells push CCT front end and elongate CCTs in length behind. 7. A single piece of CCT can be up to 110 meters (m) in length in cancer patients. 8. At the end of CCT terminals, cancer cells proliferate and increase cancer cell numbers, and form cancer cell masses in CCT terminals. All cancer cell individuals secrete tiny membrane-enclosed vesicles, fuse into CCT membrane, and grow into superlarge cytocapsulae (CC), wrapping the cancer cell masses inside. These cancer cell masses (CMs) release VEGFs and many other factors, and promote angiogenesis into the CMs to provide more nutrients and oxygen. Then, all the 5 compartments integrate and form malignant tumors in vivo in patients. 9. All original/primary and secondary tumors are physically interconnected together via CCT networks and form super-structure of CNTS system. 10. CNTSs continuously disseminate to other sites across tissues and organs via superlarge CCTs and CCT networks, including bone marrow and brain.

In many cases of pharmacotherapies, conventional cancer drugs successfully shrink big tumors, but do not actually kill cancer cells in the compact tumors, and only disperse cancer cells from compact tumors into CCT networks, that later can develop into many small cancer cell masses. After several months, the small cancer cell masses will evolve and grow, and CT-scan will find tumors everywhere. These relapsed tumors will display pan-resistance to all available cancer drugs. The artificial effectiveness of cancer drug candidates in the clinical trials can lead to approval of cancer drugs that actually promote CCTs and CCT network development, which subsequently aids in cancer metastasis and relapse, accelerating cancer patient death. With the CCT membrane acting as a barrier and protecting cancer cells from conventional drugs, physicians have to increase the drug dosage, increasing the side effects to normal cells, particularly blood cells.

In summary, the aforementioned newly discovered structural compartments of cancer cells, malignant tumors, and CNTSs, significantly expand our understanding of the major parts and the nature of cancer. This expanded understanding will effectively and efficiently accelerate the process of finding a cure for all kinds of cancers.

## Discussion

The comprehensive understanding of the complete structural compartments of cancer cells, malignant tumors, and tumor systems are essential for efficient cancer diagnosis, drug development and pharmacotherapy^25-27^. Here, we systematically investigated the complete structural compartments of cancer cells, malignant tumors, and CNTSs in 307 kinds of cancers. This study discovered multiple novel structural compartments of single cancer cells, malignant tumors and CNTSs, and paves the way to cure all kinds of solid and liquid cancers.

### The complete structure of a cell

The understanding of the complete structure of a cell is fundamental for all aspects of life on earth. Here, we identified that eukaryotic cells of human cancer cells of 307 kinds of cancers generate cytocapsulae or cytocapsular tubes outside of the cell membrane, and found the 4 major structural compartments of a cancer cell: 1) cytocapsulae or cytocapsular tubes outside of the cell membrane; 2) cytocapsular lumen fluids containing many tiny, membrane-enclosed vesicles, 3) cell membrane and the cytoplasm, and 4) nuclear membrane and the nucleus. The outer membrane outside the cell membrane can be tracked back to 2.3-2.7 billion years ago in photosynthesis gram negative bacteria. The comprehensive and consistent evidences of molecular proteins, structures, biological functions and the repeatable generation of CC and CCTs in vitro and universally of cancer cells, the outer membrane of gram-negative bacteria, chloroplast and mitochondria, demonstrated that the outer membrane beyond the cell membrane contributes multiple advantages for cell survival under the stressful microenvironments. During evolution, the outer membrane degenerated in most of cells and species on earth, and only a few species (gram negative bacteria, chloroplast and mitochondria) retained it. The CC or CCT is a temporospatial compartment of devolution of the inherent ancient capacities of generation of outer membrane in the eukaryotic cell (cancer cell). The novel complete structures of cancer cells with cytocapsulae and cytocapsular lumen fluids may provide an opportunity to understand the evolution mechanisms of cell structures during the 2.7 billion years on earth. The newly found structures of human cancer cells may expand the understanding of complete structures of eukaryotic cells with the outer membranes under stressful conditions linking to diseases and therapy development.

### The complex structure of cancer in cell, tumor and system levels

The complete structure of a cancer cell is composed of 4 major compartments: 1) cytocapsulae or cytocapsular tubes; 2) cytocapsular lumen fluids; 3) cell membrane and the cytoplasm, and 4) nuclear membrane and the nucleus. The comprehensive structure of a malignant tumor in patients consists of 5 major compartments: 1) superlarge cytocapsulae enclosing cancer cell masses, 2) CCTs and CCT networks within or outside of CMs, 3) CCT network-cancer cell masses systems, 4) CCTs twining around blood and lymph vessels, 5) mixed tissues and cells. The complete structure of a CNTS in patients is made of 4 major structural compartments: 1) all malignant tumors interconnected via CCT networks; 2) all superlarge and 3D CCT networks, 3) CCTs invade into bone marrow, and 4) CCTs invade blood-brain barrier and reach the brain. These comprehensive structural compartments of single cancer cells, malignant tumors and CNTSs in cancer patient pave avenues for future cancer researches, appropriate experimentation assays for cancer drug development, effective and efficient clinical trials, precise cancer diagnosis, and for a cure of all kinds of cancers.

In order to obtain more valuable experimental results, and reach better cancer therapy outcomes, it is suggested that conventional research or clinical activities with incomplete structures of cancer cells and/or malignant tumors to be updated, such as: 1) cell mass assays without CC or CCTs *in vitro*, 2) tumor tests in animal models without CCTs and CCT networks *in vivo*, 3) drug candidates in clinical trials without effectively inhibiting CNTS, 4) cancer drugs that do not effectively inhibit CNTSs and cure cancers. The understanding of the complete structural compartments elucidated in this study will accelerate the cure of cancer.

### Cytocapsulae and cytocapsular tubes are golden targets for effective and efficient cancer diagnosis and therapy

The aforementioned comprehensive and systematic analyses suggested that: 1) the cytocapsulae (in tissues) cytocapsular tubes (CCTs, in tissues), CCT degraded strands (thick, thin, very thin; in tissues), CCT membrane fragments and debris (in intercellular fluids; possibly in blood), CCT marker proteins/peptides/DNAs (DNA fragments) (in intercellular fluids; possibly in blood), can be golden targets for measurements in precise and decisive cancer diagnosis of all kinds of solid and liquid cancers in the pre-stage and early stage for early cancer diagnosis, before and after surgery, treatments and therapies for effects evaluation, monitoring and guidance for the treatment steps; 2) important CC or CCT marker genes or proteins can be molecular golden targets for cancer drug discovery and development aimed for effectively inhibiting and eradicating the complete compartments of CNTSs, CCT networks, malignant tumors, CC wrapped cancer cell masses, and all kinds of cytocapsular membrane enclosed cancer cells; 3) the cancer cells with CC/CCT in vitro, cancer cell masses with CC/CCTs in vitro, malignant tumors (with CC, CCTs and CCT networks) in animal, and malignant systems in animals (such as CCTX), which contain the complete compartments in each level, can be potent tools and methods for highly efficient cancer drug discovery and development; 4) The conventional cancer drug evaluation assays, which are based on inhibitory effects on incomplete compartments (cancer cells without CC/CCTs, cancer cell masses without CC/CCTs, no malignant tumor inhibition data, no CNTS inhibition data) would better be replaced with updated evaluation methods with inhibition data on complete compartments of cancer cells with CC/CCTs, cancer cell masses with CC/CCTs, malignant tumor with 5 complete compartments, and CNTs in cancer patients. Thus, these selected cancer drugs will exhibit highly effective and efficient cancer inhibition and achieve satisfactory therapy outcomes.

In summary, this study comprehensively identified the complete compartments of cancer cells, malignant tumors, and tumor systems (CNTSs) and therefore pave avenues for further cancer research and cancer drug development that will lead to the cure of all kinds of cancers.

## Methods and Materials

### Tissue slides, antibodies, and reagents

Formalin-fixed paraffin-embedded (FFPE) cancer tissue microarray slides, CDX and PDX tissue slides, and blood samples were ordered from US Biomax, Charles River laboratories, and local hospitals. Monoclonal mouse anti-γ-actin antibodies were ordered from Abcam (ab123034). Polyclonal and monoclonal rabbit anti-CM-01 (code, not real protein name) antibodies were self-developed. Secondary antibodies of anti-mouse and anti-rabbit antibodies were ordered from Thermo Fisher Scientific. DAPI was ordered from VWR. Polyclonal and monoclonal rabbit anti-CH-5 and anti-CH-6 (codes, not real protein names) antibodies were self-developed. Secondary antibodies of anti-mouse and anti-rabbit antibodies were ordered from Thermo Fisher Scientific. DAPI was ordered from VWR. Matrigel matrix was ordered from Sigma or Corning. *NOD-SCID* (strain name: *NOD*.*CB17-Prkdcscid/J*) were ordered from The Jackson Laboratory or Charles River Laboratories. All mouse work were reviewed and permitted by Cytocapsula Research Institute Animal Committee and performed in Charles River Laboratories.

### Immunohistochemistry (IHC) fluorescence staining assay

The sectioned FFPE clinical cancer tissue specimens were subjected to double immunohistochemistry staining with anti–CM-01 (1:200) and anti–γ-actin (1:200) primary antibodies and DAPI (1:1,000) for 4h at 4 °C, followed by incubation with appropriate secondary antibodies for 1h at 4 °C in a dark room. Fluorescence images were taken with a Nikon 80i upright microscope with a 20× or 40× lens. All images were obtained using MetaMorph image acquisition software and were analyzed with ImageJ software.

### Cell cytometry

Patient breast cancer cells were extracted from the fresh clinical cancer tissues, and cultured with culturing situations mimicking *in vivo* environments: 37°C, 5% CO_2_, humidity and a series of other conditions. The CH-5^high^/CH-6^high^ patient cancer cell subpopulations were isolated by fluorescence-activated cell sorting (FACS) assay with self-developed FITC-conjugated anti-CH-5 and PE-conjugated anti-CH-6 antibodies.

### Tumor xenograft formation assay

In the tumor xenografted assay, the indicated patient cancer cells, and CH-5^high^/CH-6^high^ subpopulation breast cancer cells (cell numbers: 0.5million) were mixed with 100μl Matrigel/cell culture media mixture (Matrigel: cell culture media = 1:2) (Sigma or Corning). Indicated cancer cells/Matrigel/ cell culture media mixtures were injected into *NOD/SCID* female for breast cancer xenograft (the Jackson Laboratory) by subcutaneous injection. After the tumor formation (about 75 mm^3^in volume, 5 mice/group), mice were sacrificed and tumors were excised. Tumor tissue samples were used for immunohistostaining and CCT analyses. The mouse experiments were performed according to the policies of Cytocapsula Research Institute Animal Committee and Charles River Laboratories.

### Statistical analysis

Quantitative data were statistically analyzed (mean ± SD, *t*-test, two-tailed). Statistical significance was determined by *t*-test. Significance was expressed as: *: *p*< 0.1; **: *p*< 0.05; ***: *p*< 0.01, or with the *p*-value. *P*<0.05 was considered significant.

Supported by a grant (to Dr. Yi) from Cytocapsula Research Institute Fund for Cytocapsular Tube Conducted Cancer Metastasis Research, and a grant (to Dr. Yi) from Chalst Inc Fund for Cytocapsular Tube Cancer Metastasis Research.

We greatly acknowledge Dr. Ed Harlow of Harvard Medical School (USA) and Cancer Institute of University of Cambridge (UK), and Dr. Nahum Sonenberg of McGill University of Canada, Dr. Yubo Yang, Dr. Qiping Hou and Mr. James Ranieri for their help and meaningful discussion in the study.

## Supplementary materials

**Fig. S1.**
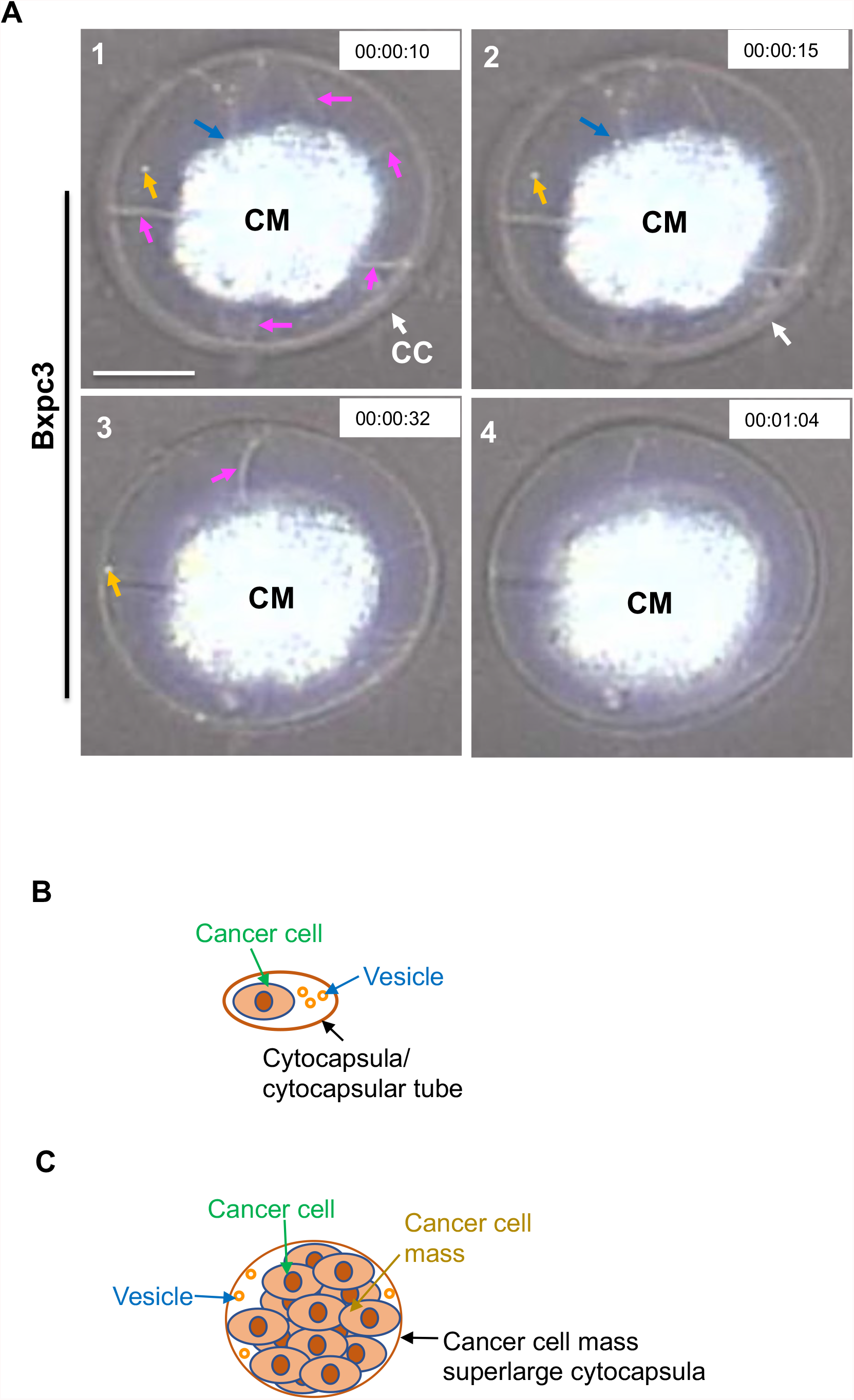
Mechanics of cancer cell mass superlarge cytocapsulae (CC) growth up. **(A)** There is a pancreas cancer Bxpc3 cell mass (CM)in the 3D Matrigel matrix layer. The Bxpc3 CM has generated a superlarge cytocapsula (CC, white arrows) outside, wrapping the CM inside. There are multiple spike-like bars (purple arrows) surrounding the cell mass and linking the cell mass and the CC membranes, supporting the superlarge CC membranes and maintain the CC morphology and shape. In the superlarge lumen and outside of the cell mass, there are hundreds of tiny, membrane-enclosed vesicles, which are in constant and random moving. The orange arrows in panels 1, 2 and 3 show a tiny, membrane-enclosed vesicle that moves into the superlarge CC membrane, and later fuse into the superlarge CC membrane and disappeared in panel 4. The blue arrows in panels 1 and 2 show a tiny, membrane-enclosed vesicle secreted by a single cancer cell in the cancer cell mass (in panel 1) that then leaves the cell mass (in panel 2). Time in the movie is shown. **(B)** A diagram of the complete compartments of a cancer cell in cancer patient tissues and organs. **(C)** A diagram of the complete compartments of a cancer cell mass in cancer patient tissues and organs.

**Fig. S2.**
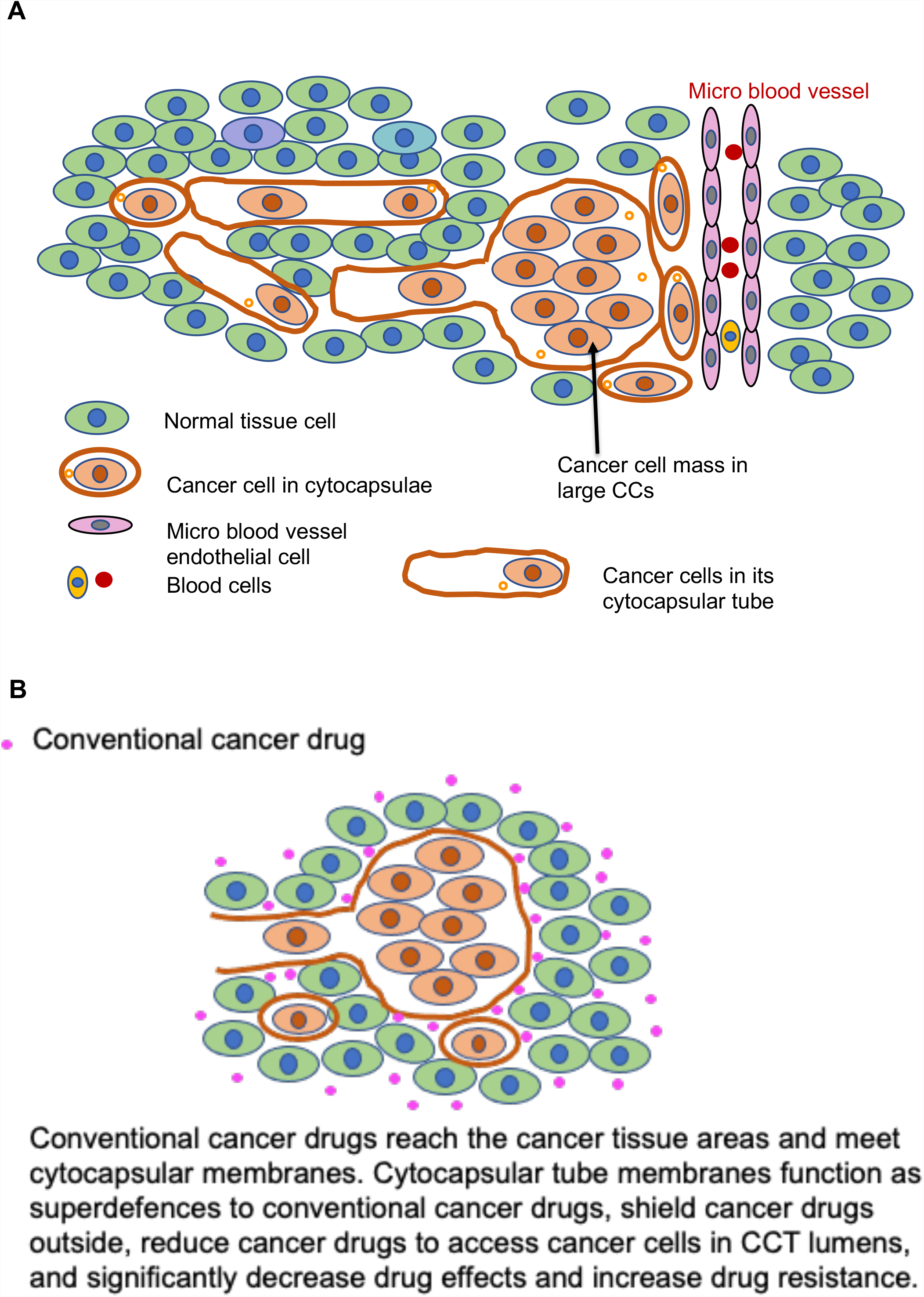
Diagram drawings of the complete compartments of malignant tumors and mechanisms of conventional cancer drug resistance of malignant tumors. (A) A diagram of the complete compartments of malignant tumors in cancer patient bodies. (B) A diagram of mechanisms of conventional cancer drug resistance of malignant tumors. The superlarge cytocapsular membrane shields most of conventional cancer drugs outside, reduces drugs to penetrate into CC lumens, decreases cancer drugs to access cancer cells in cancer cell masses, and exhibits increased pan-resistance to conventional cancer drugs, leading to weak or poor cancer therapy outcomes.

**Table S1.** The 28 subtypes of investigated blood cancers in bone marrow in Table 1.

| Cancer types | Major cancer subtypes |  | Cancer subtypes |  |
| --- | --- | --- | --- | --- |
| Blood | 1 | Acute myeloid leukemia (AML) | 1 | Acute promyelocytic leukemia (APL) |
|  |  |  | 2 | Acute myeloblastic leukemia with maturation |
|  |  |  | 3 | Acute myelomonocytic leukemia |
|  |  |  | 4 | Acute myelomonocytic leukemia with eosinophilia |
|  |  |  | 5 | Acute monocytic leukemia |
|  |  |  | 6 | Acute erythroid leukemia |
|  |  |  | 7 | Acute megakaryoblastic leukemia |
|  | 2 | Acute lymphocytic leukemia (ALL) | 1 | Acute Lymphoblastic Leukemia. ALL Subtypes. Childhood ALL. |
|  |  |  | 2 | Acute Myeloid Leukemia. |
|  |  |  | 3 | Chronic Lymphocytic Leukemia. |
|  |  |  | 4 | Chronic Myeloid Leukemia. |
|  |  |  | 5 | Hairy Cell Leukemia. |
|  |  |  | 6 | Chronic Myelomonocytic Leukemia. |
|  |  |  | 7 | Juvenile Myelomonocytic Leukemia. |
|  |  |  | 8 | Large Granular Lymphocytic Leukemia. |
|  |  |  | 9 | Childhood AML |
|  |  |  | 10 | Chronic Lymphocytic Leukemia |
|  |  |  | 11 | Chronic Myeloid Leukemia |
|  |  |  | 12 | Hairy Cell Leukemia |
|  |  |  | 13 | Chronic Myelomonocytic Leukemia |
|  |  |  | 14 | Juvenile Myelomonocytic Leukemia |
|  |  |  | 15 | Large Granular Lymphocytic Leukemia |
|  |  |  | 16 | Blastic Plasmacytoid Dendritic Cell Neoplasm |
|  |  |  | 17 | B-Cell Prolymphocytic Leukemia (B-PLL) |
|  |  |  | 18 | T-Cell Prolymphocytic Leukemia (T-PLL) |
|  | 3 | Chronic Myeloid leukemia (CML) | 1 | CML |
|  | 4 | Chronic lymphocytic leukemia (CLL) | 1 | B-cell CLL |
|  |  |  | 2 | T-cell prolymphocytic leukemia |

**Table S2.**
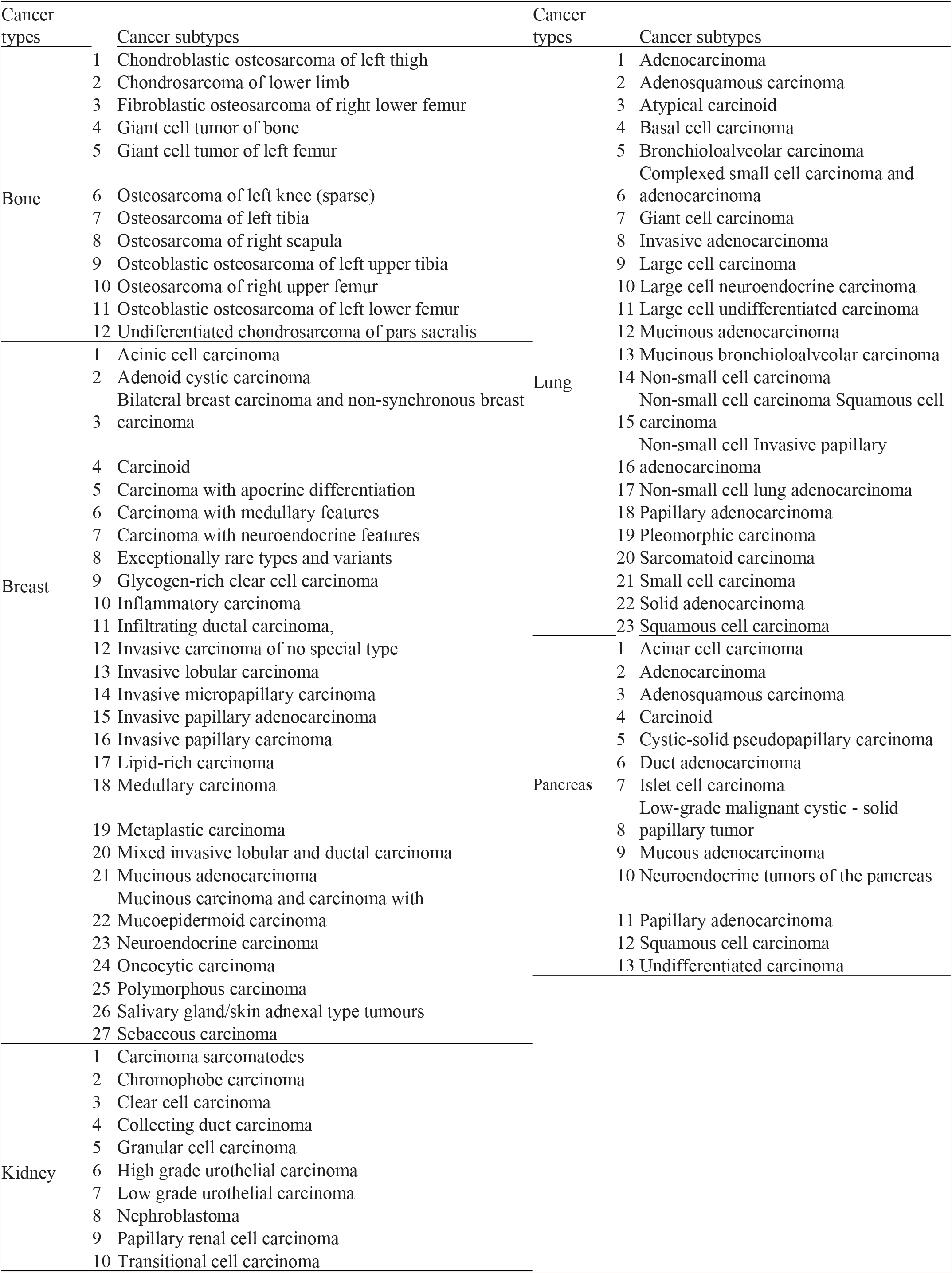
Subtypes of solid cancers with 10 (or more) kinds of investigated subtypes in Table 1.

**Table S3.**
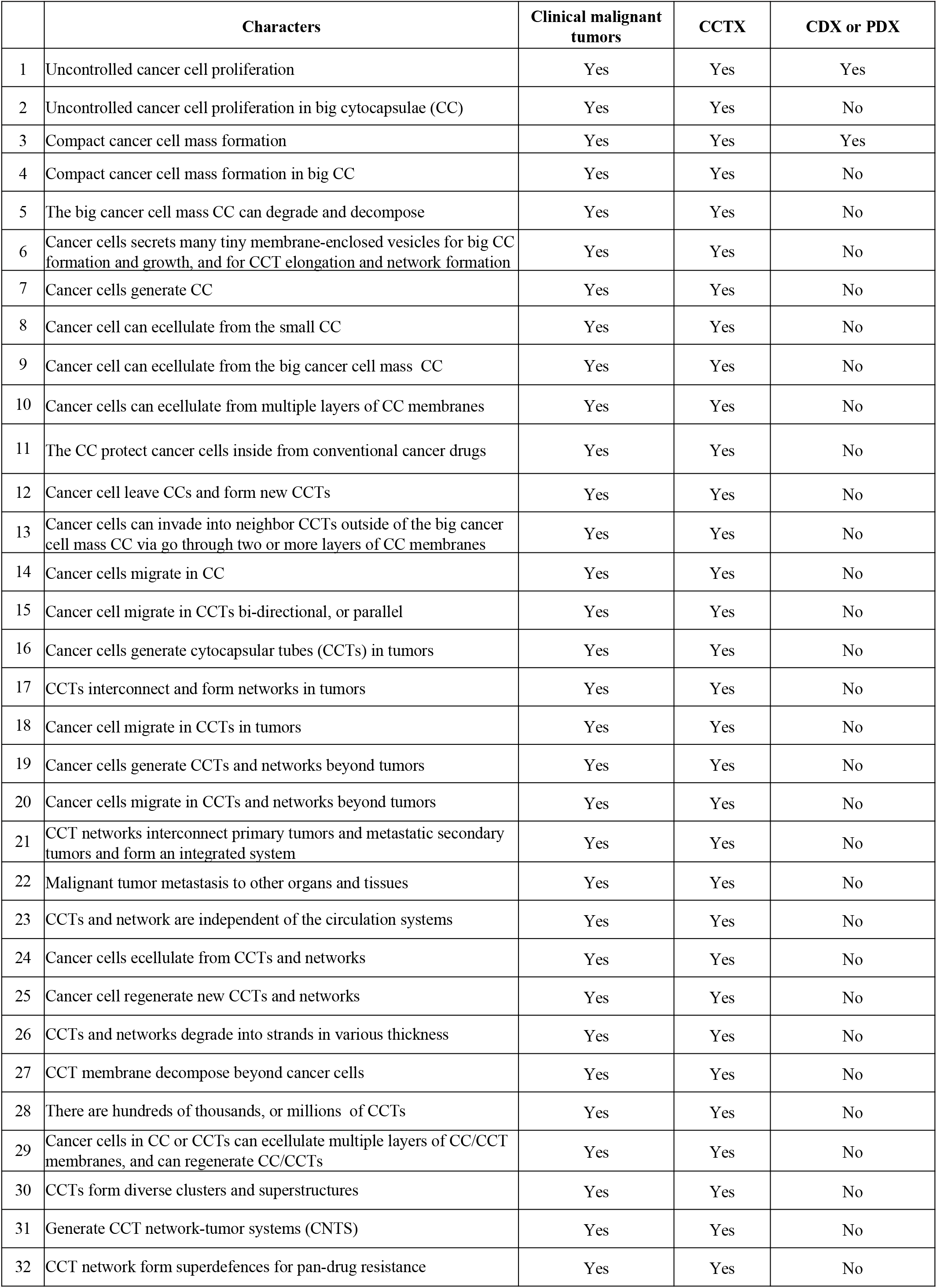
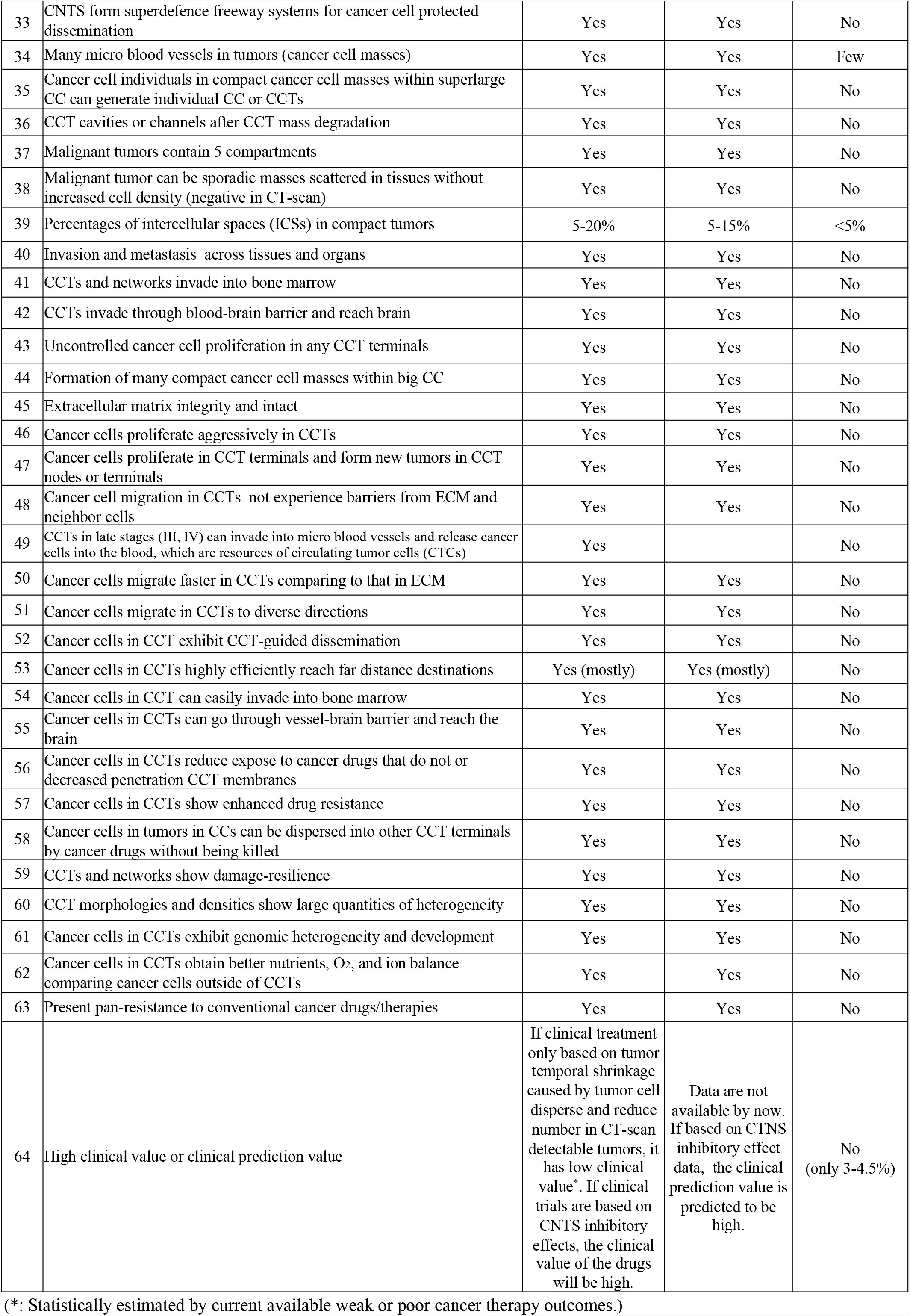
Comprehensive similarities and differences between clinical malignant tumors, CCTX and animal transplantation cancer cell masses (such as in cancer cell derived xenograft (CDX) or patient cancer cell derived xenograft (PDX)).

|  | Characters | Clinical malignant tumors | CCTX | CDX or PDX |
| --- | --- | --- | --- | --- |
| 1 | Uncontrolled cancer cell proliferation | Yes | Yes | Yes |
| 2 | Uncontrolled cancer cell proliferation in big cytocapsulae (CC) | Yes | Yes | No |
| 3 | Compact cancer cell mass formation | Yes | Yes | Yes |
| 4 | Compact cancer cell mass formation in big CC | Yes | Yes | No |
| 5 | The big cancer cell mass CC can degrade and decompose | Yes | Yes | No |
| 6 | Cancer cells secrete many tiny membrane-enclosed vesicles for big CC formation and growth, and for CCT elongation and network formation | Yes | Yes | No |
| 7 | Cancer cells generate CC | Yes | Yes | No |
| 8 | Cancer cell can ecellulate from the small CC | Yes | Yes | No |
| 9 | Cancer cell can ecellulate from the big cancer cell mass CC | Yes | Yes | No |
| 10 | Cancer cells can ecellulate from multiple layers of CC membranes | Yes | Yes | No |
| 11 | The CC protect cancer cells inside from conventional cancer drugs | Yes | Yes | No |
| 12 | Cancer cell leave CCs and form new CCTs | Yes | Yes | No |
| 13 | Cancer cells can invade into neighbor CCTs outside of the big cancer cell mass CC via go through two or more layers of CC membranes | Yes | Yes | No |
| 14 | Cancer cells migrate in CC | Yes | Yes | No |
| 15 | Cancer cell migrate in CCTs bi-directional, or parallel | Yes | Yes | No |
| 16 | Cancer cells generate cytocapsular tubes (CCTs) in tumors | Yes | Yes | No |
| 17 | CCTs interconnect and form networks in tumors | Yes | Yes | No |
| 18 | Cancer cell migrate in CCTs in tumors | Yes | Yes | No |
| 19 | Cancer cells generate CCTs and networks beyond tumors | Yes | Yes | No |
| 20 | Cancer cells migrate in CCTs and networks beyond tumors | Yes | Yes | No |
| 21 | CCT networks interconnect primary tumors and metastatic secondary tumors and form an integrated system | Yes | Yes | No |
| 22 | Malignant tumor metastasis to other organs and tissues | Yes | Yes | No |
| 23 | CCTs and network are independent of the circulation systems | Yes | Yes | No |
| 24 | Cancer cells ecellulate from CCTs and networks | Yes | Yes | No |
| 25 | Cancer cell regenerate new CCTs and networks | Yes | Yes | No |
| 26 | CCTs and networks degrade into strands in various thickness | Yes | Yes | No |
| 27 | CCT membrane decompose beyond cancer cells | Yes | Yes | No |
| 28 | There are hundreds of thousands, or millions of CCTs | Yes | Yes | No |
| 29 | Cancer cells in CC or CCTs can ecellulate multiple layers of CC/CCT membranes, and can regenerate CC/CCTs | Yes | Yes | No |
| 30 | CCTs form diverse clusters and superstructures | Yes | Yes | No |
| 31 | Generate CCT network-tumor systems (CNTS) | Yes | Yes | No |
| 32 | CCT network form superdefences for pan-drug resistance | Yes | Yes | No |
| 33 | CNTS form superdefence freeway systems for cancer cell protected dissemination | Yes | Yes | No |
| 34 | Many micro blood vessels in tumors (cancer cell masses) | Yes | Yes | Few |
| 35 | Cancer cell individuals in compact cancer cell masses within superlarge CC can generate individual CC or CCTs | Yes | Yes | No |
| 36 | CCT cavities or channels after CCT mass degradation | Yes | Yes | No |
| 37 | Malignant tumors contain 5 compartments | Yes | Yes | No |
| 38 | Malignant tumor can be sporadic masses scattered in tissues without increased cell density (negative in CT-scan) | Yes | Yes | No |
| 39 | Percentages of intercellular spaces (ICSs) in compact tumors | 5-20% | 5-15% | <5% |
| 40 | Invasion and metastasis across tissues and organs | Yes | Yes | No |
| 41 | CCTs and networks invade into bone marrow | Yes | Yes | No |
| 42 | CCTs invade through blood-brain barrier and reach brain | Yes | Yes | No |
| 43 | Uncontrolled cancer cell proliferation in any CCT terminals | Yes | Yes | No |
| 44 | Formation of many compact cancer cell masses within big CC | Yes | Yes | No |
| 45 | Extracellular matrix integrity and intact | Yes | Yes | No |
| 46 | Cancer cells proliferate aggressively in CCTs | Yes | Yes | No |
| 47 | Cancer cells proliferate in CCT terminals and form new tumors in CCT nodes or terminals | Yes | Yes | No |
| 48 | Cancer cell migration in CCTs not experience barriers from ECM and neighbor cells | Yes | Yes | No |
| 49 | CCTs in late stages (III, IV) can invade into micro blood vessels and release cancer cells into the blood, which are resources of circulating tumor cells (CTCs) | Yes |  | No |
| 50 | Cancer cells migrate faster in CCTs comparing to that in ECM | Yes | Yes | No |
| 51 | Cancer cells migrate in CCTs to diverse directions | Yes | Yes | No |
| 52 | Cancer cells in CCT exhibit CCT-guided dissemination | Yes | Yes | No |
| 53 | Cancer cells in CCTs highly efficiently reach far distance destinations | Yes (mostly) | Yes (mostly) | No |
| 54 | Cancer cells in CCT can easily invade into bone marrow | Yes | Yes | No |
| 55 | Cancer cells in CCTs can go through vessel-brain barrier and reach the brain | Yes | Yes | No |
| 56 | Cancer cells in CCTs reduce expose to cancer drugs that do not or decreased penetration CCT membranes | Yes | Yes | No |
| 57 | Cancer cells in CCTs show enhanced drug resistance | Yes | Yes | No |
| 58 | Cancer cells in tumors in CCs can be dispersed into other CCT terminals by cancer drugs without being killed | Yes | Yes | No |
| 59 | CCTs and networks show damage-resilience | Yes | Yes | No |
| 60 | CCT morphologies and densities show large quantities of heterogeneity | Yes | Yes | No |
| 61 | Cancer cells in CCTs exhibit genomic heterogeneity and development | Yes | Yes | No |
| 62 | Cancer cells in CCTs obtain better nutrients, O<sub>2</sub>, and ion balance comparing cancer cells outside of CCTs | Yes | Yes | No |
| 63 | Present pan-resistance to conventional cancer drugs/therapies | Yes | Yes | No |
| 64 | High clinical value or clinical prediction value | If clinical treatment only based on tumor temporal shrinkage caused by tumor cell disperse and reduce number in CT-scan detectable tumors, it has low clinical value*. If clinical trials are based on CNTS inhibitory effects, the clinical value of the drugs will be high. | Data are not available by now. If based on CTNS inhibitory effect data, the clinical prediction value is predicted to be high. | No<br>(only 3-4.5%) |

(\*: Statistically estimated by current available weak or poor cancer therapy outcomes.)

